# Children’s syntax is supported by the maturation of BA44 at 4 years, but of the posterior STS at 3 years of age

**DOI:** 10.1101/2022.09.17.508373

**Authors:** Cheslie C. Klein, Philipp Berger, Tomás Goucha, Angela D. Friederici, Charlotte Grosse Wiesmann

**Affiliations:** Department of Neuropsychology, Max Planck Institute for Human Cognitive and Brain Sciences, Leipzig, Germany; Research Group Milestones of Early Cognitive Development, Max Planck Institute for Human Cognitive and Brain Sciences, Leipzig, Germany

**Keywords:** Broca’s area, Cortical maturation, Early childhood, Language development, Syntax development

## Abstract

Within the first years of life, children learn major aspects of their native language. However, the ability to process complex sentence structures, a core faculty in human language called syntax, has been found to emerge only slowly. A milestone in the acquisition of syntax is reached around the age of 4, when children learn a variety of syntactic concepts, including, for example, subordinate clauses. Here, we ask which maturational changes in the child’s brain underlie the emergence of syntactic abilities around this critical age. We relate markers of cortical brain maturation to 3- and 4-year-olds’ syntactic in contrast to other language abilities. Our results show that distinct cortical brain areas support syntax in the two age groups: While 3-year-old children’s syntactic abilities were associated with increased surface area in the most posterior part of the left superior temporal sulcus, 4-year-old children showed an association with cortical thickness in the left posterior part of Broca’s area, i.e. BA44. The present findings suggest that syntactic abilities rely on the maturation of distinct cortical regions in 3- compared to 4-year-olds. The observed shift to more mature regions involved in syntax may underlie the behavioral milestones in syntax acquisition around 4 years of age.

---

Language comprehension and production are among the most demanding cognitive abilities to be acquired during development. In order to gain full proficiency in their language, a child has to successfully master each of its modular components and learn to integrate them altogether. Already within the first year of life, children learn to segment the incoming speech stream to phonological word forms and to associate them with meaning stored in the mental lexicon (Speer and Ito 2009; Bergelson and Swingley 2012). Before the age of two, children can use prosodic cues for the detection of phrase boundaries and can organize words into morphosyntactic categories (Soderstrom 2003; Höhle et al. 2004). However, the ability to engage in the rule-based formation of words into a hierarchical sentence structure, what is called syntax, develops comparably slowly and is not fully mastered until young adulthood (Skeide and Friederici 2016; Friederici 2020). Syntax is a core faculty in human language and serves as the foundation for uniquely human communication, which sets it apart from other species (Friederici 2018, 2020). Parallel developmental trajectories in children’s acquisition of syntax and the emergence of other higher-order cognitive functions have been found (Milligan et al. 2007; Friedman and Sterling 2019), emphasizing the impact on early cognitive development. Between 3 and 4 years of age, developmental milestones are observed in the ability to process and produce complex sentence structures (Fox and Grodzinsky 1998; Akhtar 1999; Tomasello and Brooks 1999; Guasti 2002; Kauschke 2012). Particularly, while at 3 years of age, children reliably process the canonical word order of their language (Akhtar 1999; Dittmar et al. 2008; Schipke et al. 2012), more complex syntactic phrases, such as long-distance wh-questions, passive constructions and subordinate clauses, are not fully acquired before the fourth birthday (Fox and Grodzinsky 1998; Tomasello and Brooks 1999; Guasti 2002; Kauschke 2012). What are the neural maturational processes that underlie this critical step in human language development? In the present study, we set out to investigate the maturation of brain structure in the critical age range between 3 and 4 years that support the acquisition of complex syntax.

In adults, language processing has been found to demand multiple brain areas to interact, together forming what has been referred to as the language network in the human brain (Friederici 2002). The core of this network includes Broca’s area, which consists of the pars opercularis (BA44) and the pars triangularis (BA45) in the left inferior frontal gyrus (IFG; Skeide and Friederici 2016), and Wernicke’s area in the superior temporal lobe (DeWitt and Rauschecker 2013). In the mature brain, BA45 in the anterior part of the IFG is functionally involved in the processing of lexical meaning (semantics) together with the temporal lobe. Conversely, BA44 in the posterior part of the IFG is involved in the integration of complex syntax in a multitude of studies (Friederici et al. 2000, 2006; Friederici 2002, 2011; Makuuchi et al. 2009; Newman et al. 2010; Goucha and Friederici 2015; Zaccarella et al. 2015). For example, Goucha and Friederici (2015) reported an activation in both parts of the left IFG when adult participants heard sentences consisting of words without meaning but with correct syntactic order and derivational and inflectional morphological elements. After removing the meaning-conveying derivational morphological elements leaving only elements that convey structural cues, the authors found an effect solely in BA44 suggesting that this area is crucial for syntactic processing.

Although adults process syntactic and semantic features separately (Jackendoff 2007) and, consequently, show dissociable functional representations in the brain (Skeide and Friederici 2016), behavioral and imaging studies in children indicate that this functional segregation is not completed until late childhood (Friederici 1983; Brauer and Friederici 2007; Nuñez et al. 2011; Knoll et al. 2012; Skeide et al. 2014; Wu et al. 2016; Xiao et al. 2016; Strotseva-Feinschmidt et al. 2019). Specifically, it was found that children at least until the age of 7 strongly rely on word meaning to successfully process complex sentences (Dittmar et al. 2008;

Skeide et al. 2014; Skeide and Friederici 2016). For example, while 3-year-old children have difficulties processing object-initial sentences when both the agent and patient are animate (Schipke et al. 2012), they perform above chance when only the agent of the sentence is animate, providing them with a semantic cue to resolve the verb argument structure (Strotseva-Feinschmidt et al. 2019). Skeide et al. (2014) found that children’s reliance on semantics when processing syntactically complex sentences was reflected in reduced functional segregation in the language network: children aged 3.9 to 4.11 years did not show separate effects of syntactic complexity or semantic plausibility in their neural activation, instead they exhibited an interaction of syntax and semantics in the posterior part of the left superior temporal lobe. The neural activation in the left BA44 for syntactic processing, found in adults, increased between 4 and 9 years of age (Skeide et al. 2016). An increase in activation in the left IFG with increasing syntactic proficiency from 5 years of age was confirmed by a number of further functional brain imaging studies that also underline that the interference of syntax and semantics is reflected in an immature brain activation pattern (Brauer and Friederici 2007; Nuñez et al. 2011; Knoll et al. 2012; Wu et al. 2016; Xiao et al. 2016). In sum, on a brain functional level, these findings show an interaction between syntactic and semantic processes in young children with an increasing spatial segregation during middle childhood. At least by the age of 4, a gradual development toward a more adult-like pattern for syntax, with increasing activation of left inferior frontal brain regions, is initiated (Skeide and Friederici 2016). Against the background of a major behavioral improvement in complex syntax processing and production, a central question is to what extent the maturation of the left IFG contributes to the change in syntactic processing abilities observed between 3 and 4 years. In the present study, we therefore set out to investigate which brain structural changes underlie and precede the increasing functional involvement of the left IFG observed after 4 years of age.

It is known that during brain maturation in early childhood, the human cerebral cortex undergoes complex cytoarchitectonic changes, caused by the proliferation of dendrites, synaptic pruning and myelination (Walhovd et al. 2016). Using structural magnetic resonance imaging (sMRI) as an in vivo reconstruction method, cortical thickness and surface area have been suggested as indices of cortical maturation (Fischl and Dale 2000), providing spatial information about the developmental status of brain regions and networks. Cortical surface area has been shown to expand dramatically across the entire cortex between infancy and adulthood with a peak in late childhood (Li et al. 2014; Lyall et al. 2015). On a regional level, this increase in cortical surface area has been found to be positively related to cognitive development in childhood (Walhovd et al. 2016; Cafiero et al. 2019; Grosse Wiesmann et al. 2020). The developmental trajectory of cortical thickness, in contrast, is less clear, as its peak depends on complementary microphysiological processes and is subject to considerable local and interindividual variation (Shaw et al. 2006; Cafiero et al. 2019; Natu et al. 2019; Grosse Wiesmann et al. 2020). A range of studies suggest that cortical thickness generally peaks around late childhood and puberty and thins thereafter, with association cortices showing a later peak than sensory and motor cortices (Sowell 2004; Shaw et al. 2008). Accordingly, depending on the age, cognitive domain, and brain region, cortical thickness has been found to be either positively or negatively related to cognitive function (Shaw et al. 2006; Cafiero et al. 2019; Grosse Wiesmann et al. 2020).

In early preschool age, the relation of cortical brain structure and cognitive function has rarely been studied to date, and, in particular, research on the relation to early language and syntactic development is lacking. In children aged 4 years and older, some evidence exists that language development is related to cortical maturation in the language network, and particularly the left IFG (Sowell 2004; Lu et al. 2006; Richardson et al. 2010; Fengler et al. 2015; Qi et al. 2019, 2021). In addition to a relation of children’s sentence comprehension abilities with gray matter indices in the left IFG, Fengler et al. (2015) also found an association in the inferior parietal lobe and posterior superior temporal gyrus from 5 years of age. While several of these studies have investigated brain structural changes involved in sentence comprehension from 4 or 5 years and older (Fengler et al. 2015; Qi et al. 2019, 2021), the maturational changes of those brain structures that support early syntax acquisition before the age of 4 years, when children start to master more complex sentence structures, remain largely unknown.

This leaves a gap in the literature on the brain maturational changes underlying the important syntactic milestones achieved between 3 and 4 years of age. Further, neuroimaging studies assessed children’s knowledge of the syntactic rule system in sentence comprehension tasks (Brauer and Friederici 2007; Nuñez et al. 2011; Knoll et al. 2012; Skeide et al. 2014, 2016; Fengler et al. 2015; Wu et al. 2016; Xiao et al. 2016; Qi et al. 2019), while no research on the brain maturational processes supporting early syntactic production exists so far.

In the present study, we therefore investigate the role of cortical brain maturation in early syntax development in young 3- and 4-year-old children. In particular, we focused on children’s syntax comprehension and production and compared this with their global language abilities. In a preregistered procedure, we related syntax comprehension and production as well as global language abilities extracted from a standardized test of children’s general language development to gray matter maturation indices in regions of the left-lateralized language network. We hypothesized that language performance, in general, would be associated with maturation indices in the entire left language-related brain network (Friederici and Gierhan 2013; Fedorenko and Thompson-Schill 2014). Children’s syntactic abilities, in contrast, were expected to be specifically related to the maturation of left BA44 with an increasing relevance of this region between 3 and 4 years of age, based on previous studies in older children (Fengler et al. 2015; Skeide and Friederici 2016; Skeide et al. 2016).

## Methods

This study was preregistered at https://osf.io/g9bke.

### Participants

MRI and behavioral data of 37 typically developing 3- and 4-year-old children (17 3-year-old children, mean age = 3.31, SD = 0.18, range = 3.07 to 3.59, 10 female; 20 4-year-old children, mean age = 4.31, SD = 0.17, range = 4.02 to 4.58, 11 female) were analyzed for this study. Behavioral data from a standardized test battery of general language development (SETK 3-5; *Sprachentwicklungstest für drei- bis fünfjährige Kinder: SETK3–5;* Grimm 2001) were available for a total of 60 monolingual German-speaking children, with 27 3-year-olds and 33 4-year-olds. Of these, we excluded children that (1) had an indication for a speech development disorder (n = 2; T-value < 35) in the language test, (2) did not participate in or aborted the MRI (n = 9), (3) showed motion artifacts in the sMRI data, detected by visual inspection (n = 11; Grosse Wiesmann et al. 2020), or (4) had a coincidental neurological diagnosis (n = 1). Two children did not receive all language scores, as one child did not complete all subtests of the SETK 3-5 and the transcript of one subtest of another child had been lost. Handedness was evaluated with the German version of the Edinburgh Handedness Inventory (Oldfield 1971). Parental informed consent was obtained for all children, and the study was approved by the Ethics Committee at the Faculty of Medicine of the University of Leipzig (number of approval: 090/12-ff). The data have previously been analyzed with regard to other research questions (Grosse Wiesmann, Friederici, et al. 2017; Grosse Wiesmann, Schreiber, et al. 2017; Grosse Wiesmann et al. 2020; Berger et al. preprint).

### Behavioral data

#### Global language abilities

Children underwent a standardized test battery of general language development, designed for 3- to 5-year-old children (SETK 3-5; Grimm 2001). The SETK 3-5 is a comprehensive diagnostic tool that provides the possibility to compare individual test performance to detailed age-related norms for clinical application. In the current study, we were interested in relating individual developmental differences with brain structure. To capture these individual differences, we used raw values, rather than normalized T-scores, of each subtest of the SETK 3-5 and performed z-transformation within groups of 3-year-olds and 4-year-olds, respectively, to account for differences in the item structure of the test version for 3- and for 4-year-olds. The standardization was carried out within the age groups of the full sample (N = 60). The z-scores were combined to scales for language comprehension, production and language memory, following the standard procedures in the SETK 3-5. In the last step, these scale values were aggregated to a global language score for each subject. The global language scores of 3- and 4-year-olds were normally distributed, as indicated by a Shapiro-Wilk test (3-year-olds: p = 0.24, 4-year-olds: p = 0.98), and no outliers were identified.

#### Syntactic abilities

In order to assess syntactic abilities in the same children, we made use of the subtests of the SETK 3-5 that tested sentence comprehension and production in the 3- and 4-year-olds respectively (see details in Supplementary Material). Children’s syntactic performance on the comprehension level was tested in an object manipulation task (‘Verstehen von Sätzen’; Grimm 2001). The presented sentences increase in their grammatical complexity to evaluate children’s syntactic knowledge (see details in Supplementary Material). We z-transformed raw scores within age groups to form the syntactic comprehension score. Syntactic comprehension scores were normally distributed, as indicated by a Shapiro-Wilk test (3-year-olds: p = 0.069, 4-year-olds: p = 0.13), and no outliers were identified. To assess children’s syntactic production abilities, we developed and preregistered a coding procedure that quantified the syntactic complexity of children’s produced sentences. This was achieved by identifying the length of the longest syntactically correct fragment in words per item, a proxy for syntactic production ability modified for children’s data (see Hunt 1970; Loban 1976; Nippold et al. 2014, 2017). Incomplete clauses with correct syntactic structure were included in the analysis (Vasilyeva et al. 2008). Hesitation and self-corrections were not considered as part of the fragment, but nonwords with clear part-of-speech assignment and grammatically correct inflection were treated as real words. Two immediately following adjectives were counted as one to ensure that the length of the fragment would not be extended without adding syntactic complexity to it. The length of the fragments was computed based on production data collected in two standardized subtests of the SETK 3-5 that were designed to elicit syntactically complex sentences (Grimm 2001). For the 3-year-olds, sentence production was elicited in a picture description task in which children were expected to produce prepositional phrases (‘Enkodierung semantischer Relationen’, ESR; Grimm 2001). For the 4-year-olds, sentence production was elicited in a sentence repetition task in which children were asked to reproduce sentences with correct morphosyntactic inflection and either plausible or implausible meaning (‘Satzgedächtnis’, SG; Grimm 2001). Importantly, our coding procedure was independent of the scoring system of the SETK 3-5 for these production subtests and was applied identically to both age groups. The raw scores were z-transformed within the groups of 3- and 4-year-olds to account for differences in length of the syntactic fragment that may have arisen from differences in the elicitation context. This way, the length of 3-year-olds’ syntactically correct fragments was evaluated only in relation to the length of other 3-year-olds’ fragments, whereas 4-year-olds’ fragments length were evaluated in relation to those of other 4-year-olds. These standardized values formed the syntactic production score. Testing sessions were audio recorded and language production data were transcribed. In addition, more than 20% of the audio files were transcribed by a second independent rater. Items that received conflicting transcriptions were transcribed by a third independent rater and the transcription of the majority was used. For the subtest ESR, the first and second transcription agreed in over 87% and for the subtest SG in over 93% of the sentences. A Shapiro-Wilk test revealed that syntactic production scores were normally distributed (3-year-olds: p = 0.50, 4-year-olds: p = 0.22) and had no statistical outliers. Syntactic comprehension and production scores were used as our main measures for children’s syntactic ability. An additional measure for syntax was obtained from the word production subtest (‘Morphologische Regelbildung’; Grimm 2001) in which children’s ability to assign the plural on nouns was tested to estimate their knowledge of the morpho-syntactic rule system, which is fundamental to form grammatically correct sentences (Dittmar et al. 2008). The resulting morpho-syntactic production scores were normally distributed (3-year-olds: p = 0.068, 4-year-olds: p = 0.94) with no statistical outliers.

#### General Cognitive Development

Children’s non-verbal IQ was assessed using the Kaufman Assessment Battery for Children (Kaufman ABC; Melchers and Preuß 2003), testing for visual and auditory working memory, selective attention as well as spatial representation abilities.

### MRI data acquisition

MRI data were obtained on a 3-T Siemens scanner (Siemens MRT Trio series) equipped with a 32-channel head coil. High-resolution 3D T_1_-weighted sMRI data was acquired with the MP2RAGE sequence (Marques et al. 2010) at 1.2×1×1mm resolution with the following imaging parameters: inversion time TI_1_ = 700ms; flip angle *α*_1_ = 4°; TI_2_ = 2,500ms; *α*_2_ = 5°; repetition time TR = 5,000ms; echo time TE = 3,24ms; FoV = 192 × 192mm; 176 sagittal slices; GRAPPA 3; partial Fourier phase factor 6/8; bandwidth 240HzPx; acquisition time 5:22 min (as described in Grosse Wiesmann et al. 2020).

### MRI data analysis

#### Cortical surface-based analyses

We used the preprocessed brain images from recent studies using the current dataset to obtain measures of cortical thickness and surface area (see for details Grosse Wiesmann et al. 2020; Berger et al. preprint). In brief, individual brain images were preprocessed in FreeSurfer 5.3.0 (http://surfer.nmr.mgh.harvard.edu/) following the standard surface-based pipeline (Fischl and Dale 2000) to reconstruct cortical surfaces and generate local estimates of cortical thickness and surface area. Cortical thickness, defined as the closest distance from the grey matter/white matter boundary to the grey matter/cerebrospinal fluid boundary and surface area of the grey matter/white matter boundary was calculated at each vertex (Fischl and Dale 2000). The resulting maps for cortical thickness and surface area were smoothed on the tessellated surfaces using a 10-mm FWHM Gaussian kernel. A common group template was created from the individual T_1_-weighted images of the children who participated in the study using ANTs (Avants et al. 2008). To allow for an accurate matching of local cortical thickness and surface area measures across participants, the individual cortical surfaces were registered to the common group template.

#### ROI selection

Based on previous literature (Friederici and Gierhan 2013; Fedorenko and Thompson-Schill 2014; Qi et al. 2019) and following our preregistered hypotheses (https://osf.io/g9bke), we selected language-related ROIs in the fronto-temporal cortex of the left hemisphere, using the Desikan-Killiany atlas (Desikan et al. 2006). From these, we created a language network mask, including the following areas (see Supplementary Figure 1): superior temporal gyrus, banks of the superior temporal sulcus, Heschl’s gyrus, middle temporal gyrus, inferior parietal lobule, supramarginal gyrus, insula, caudal middle frontal gyrus and inferior frontal gyrus (pars opercularis, pars triangularis and pars orbitalis). In addition, we created a mask capturing the left BA44 that was our core hypothesized ROI for syntactic abilities (see Supplementary Figure 1). For the exploratory analysis we additionally created a mask of the left BA45. All masks were generated on the common group template.

#### Statistical analysis

The relation of our main variables for global language, syntactic comprehension and syntactic production abilities with children’s cortical thickness and surface area were estimated in general linear models (GLM) using the tool mri_glmfit implemented in FreeSurfer. Our GLMs were designed to identify regions in which children’s global language or syntactic ability was associated with the respective brain maturation marker (i.e., cortical thickness or surface area). Because of task differences between 3- and 4-year-old children based on major qualitative improvements in syntax between 3 and 4 years of age (Fox and Grodzinsky 1998; Akhtar 1999; Tomasello and Brooks 1999; Grimm 2001; Guasti 2002; Kauschke 2012), we conducted separate analyses within both age groups. Further, we tested for overlapping effects in the two age groups by performing conjunction analyses using FreeSurfer’s mri_concat (Nichols et al. 2005). In addition, following our preregistration, we report GLMs across both age groups of 3- and 4-year-olds with age group as factor (see Supplementary Results). To examine the specificity of the effects, we controlled for sex, non-verbal IQ, handedness and estimated intracranial volume (eTIV). For all models, multiple comparison correction was applied with a clusterwise correction using the FreeSurfer tool mri_glmfit-sim, specifying a cluster-forming threshold of P < 0.01, clusterwise threshold of P < 0.05 (Greve and Fischl 2018), positive relation with surface area, and bidirectional relation with cortical thickness. For the clusterwise correction, a Monte Carlo simulation with 10,000 iterations was precomputed on the group template. We performed hypothesis-driven small-volume correction in pre-specified regions of interest. In particular, following the preregistration, we ran GLMs on children’s global language, syntactic comprehension and syntactic production scores in the language network mask. In addition, GLMs on the syntactic comprehension and syntactic production scores were run within the left BA44, specifically, due to our strong a-priori hypotheses for this region being involved in syntax (Goucha and Friederici 2015). Correlation coefficients for the relation between maturation indices and language scores were computed in RStudio (RStudio Team 2020). For this, each child’s mean cortical thickness/surface area from the significant clusters were extracted with FreeSurfer’s mris_anatomical_stats. These data were tested for normality using the Shapiro-Wilk test which revealed some deviation from the normal distribution. Consequently, Spearman’s rank correlation coefficient was computed throughout the analyses. There were no statistical outliers with respect to mean thickness or surface area.

## Results

To test whether language and syntactic abilities were related to cortical brain structure, we reconstructed cortical surface area and thickness from high-resolution anatomical MRI in the same children that participated in the behavioral task battery. We ran separate GLMs to indicate cortical brain regions that show significant correlations with the global language, syntactic comprehension and syntactic production scores, respectively, in 3- and 4-year-old children. We further tested for overlapping effects in the two age groups by performing conjunction analyses.

### Global language abilities and cortical brain structure

When performing small-volume correction in the language network mask, we observed no significant correlation of children’s global language scores with their cortical thickness or surface area in neither age group of 3- and 4-year-olds. In addition, the conjunction analysis revealed no cluster that overlapped significantly in the two age groups. The GLMs across both age groups yielded comparable results, as reported in the supplementary material, except for a cluster in the insula showing a positive correlation across 3- and 4-year-olds’ global language scores with their cortical thickness (see Supplementary Figure 2 and Supplementary Table 1).

### Syntactic abilities and cortical brain structure

#### Syntactic comprehension score

In our separate GLMs within the entire language network mask, we found no significant correlation of children’s syntactic comprehension scores with their cortical thickness or surface area in neither of the two age groups. The GLMs within the preregistered mask of BA44 yielded no significant effect for 3-year-olds’, but a positive correlation of 4-year-olds’ syntactic comprehension scores with cortical thickness in this ROI (see Figure 1a and Table 1). Further analysis confirmed that the positive correlation was only significant in the 4-year-olds (3-year-olds: *ρ* = −0.50, P = 0.039; 4-year-olds: *ρ* = 0.61, P = 0.004) and differed significantly between age groups (z = 5.31, P < 0.001). The effect remained significant when controlling for sex, non-verbal IQ, handedness and eTIV.

**Figure 1:**
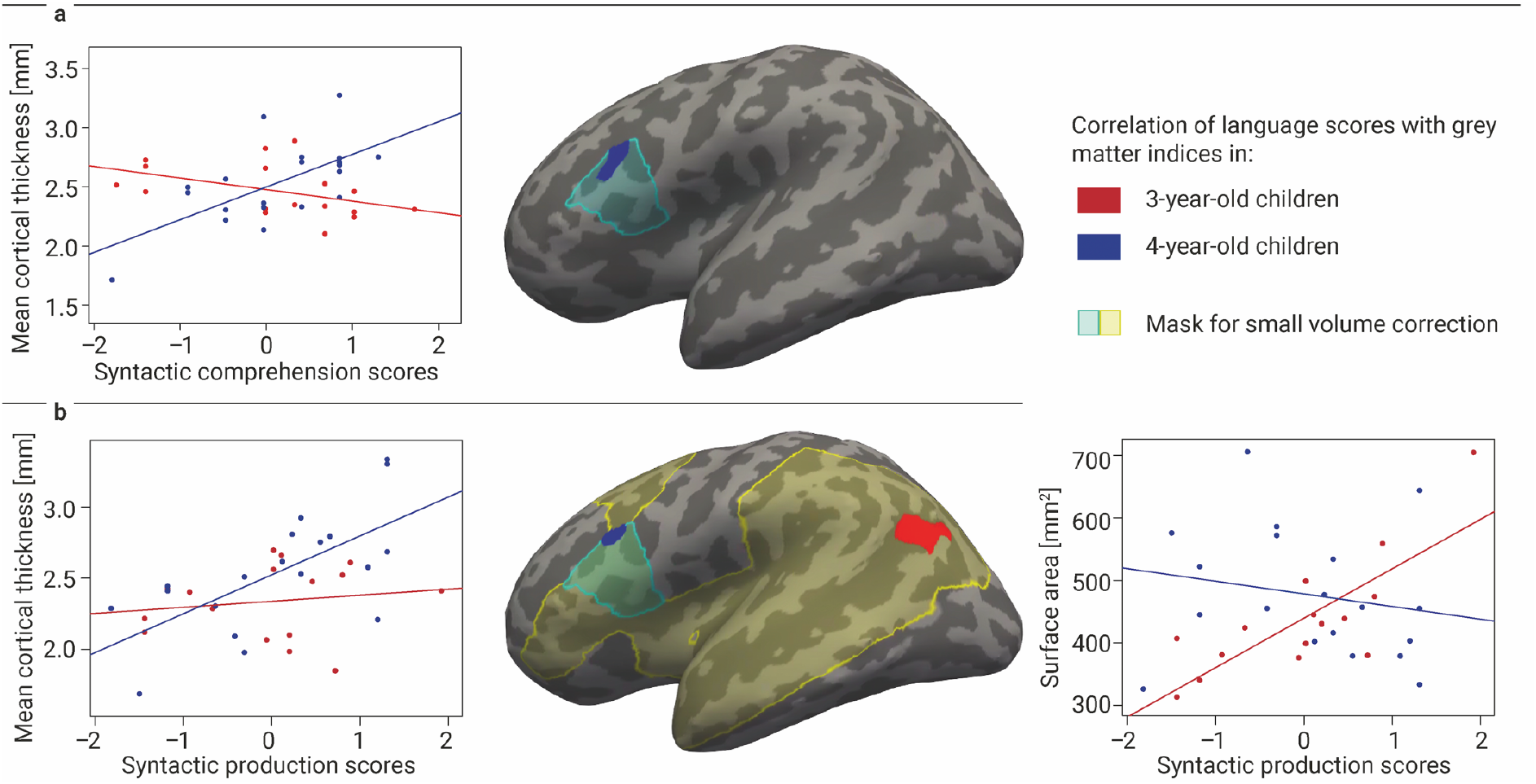
(a) Linear correlation of syntactic comprehension abilities of 4-year-old children (blue) with cortical thickness in the mask of the left BA44 (light blue). (b) Linear correlation of syntactic production abilities of children (3-year-olds: red; 4-year-olds: blue) with surface area/cortical thickness in the preregistered mask of language-related ROIs (light yellow) and of the left BA44 (light blue). The correlations were independent of sex, non-verbal IQ and eTIV. All clusters are reported at a cluster-forming threshold of P < 0.01 and clusterwise threshold of P < 0.05 and displayed on the inflated cortex of the common group template.

**Table 1:**
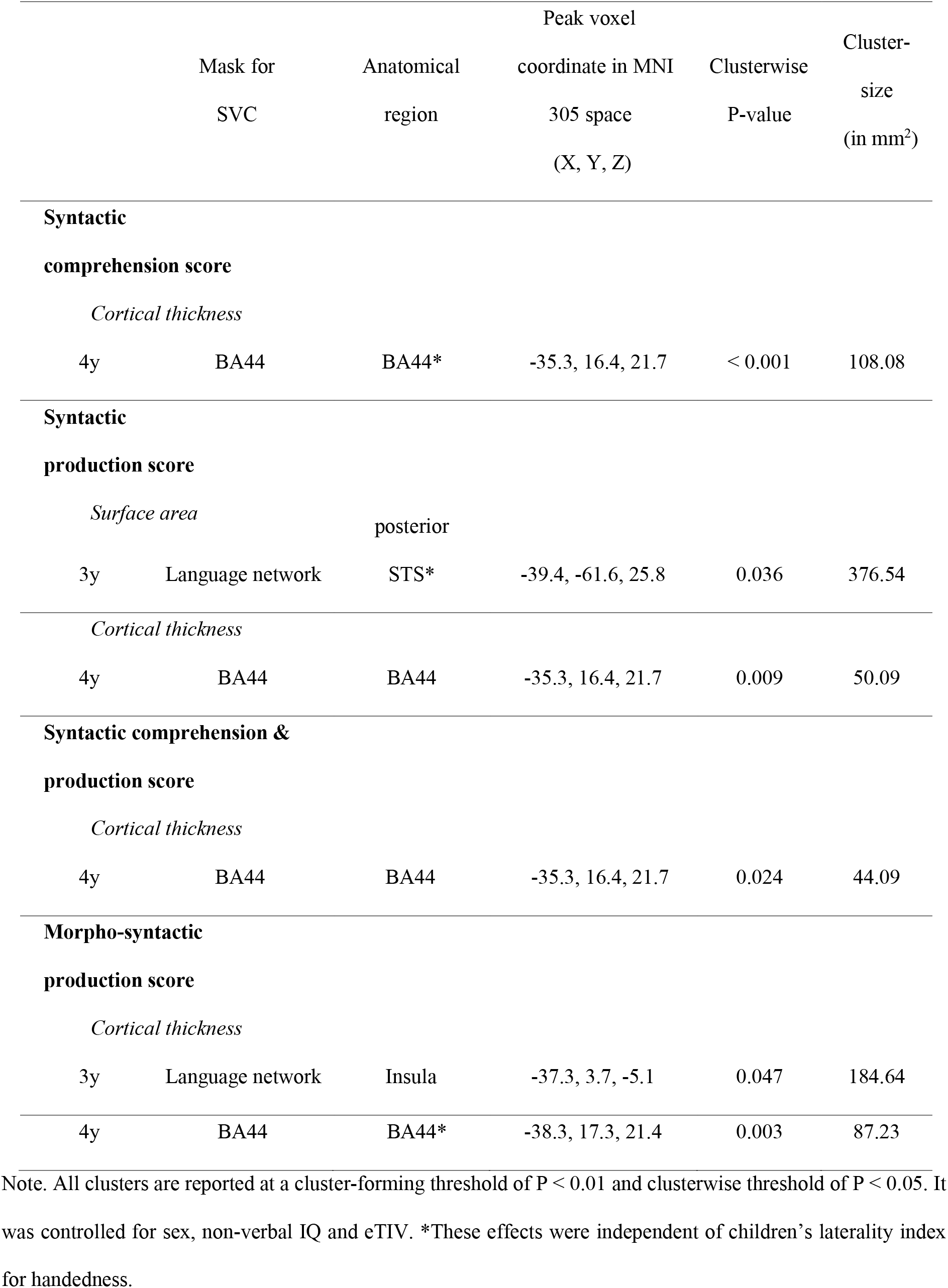
Mask for small-volume correction (SVC), anatomical region of effect, MNI coordinates, effect size, exact clusterwise P-value and cluster size of significant relations between cortical brain maturation indices and language scores in the left hemisphere.

#### Syntactic production score

When performing small-volume correction in the language network mask, 3-year-old’s but not 4-year-olds’ syntactic production scores showed a positive correlation with children’s surface area in the most posterior part of the superior temporal sulcus (STS; see Figure 1b and Table 1). This effect remained significant when controlling for sex, non-verbal IQ, handedness and eTIV. Further analysis corroborated that this relation was only present in the 3-year-olds (3-year-olds: *ρ* = 0.72, P = 0.002; 4-year-olds: *ρ* = −0.20, P = 0.40), and that the correlation coefficients in this region differed significantly between age groups (z = 4.67, P < 0.001). In the 4-year-olds, in turn, we observed a positive association of syntactic production scores with cortical thickness with a small-volume correction in BA44 (see Figure 1b and Table 1), and no effect for the 3-year-olds. Again, further analysis corroborated that this relation was only present in the 4-year-olds (3-year-olds: *ρ* = 0.17, P = 0.53; 4-year-olds: *ρ* = 0.69, P = 0.001), and that the correlation coefficients differed significantly between age groups (z = 2.79, P = 0.005). The cluster remained significant when controlling for sex, non-verbal IQ and eTIV, but regressed after adding handedness to the model (see Supplementary Material).

We found no significant conjunction between 3- and 4-year-old children’s syntactic comprehension or production score within the language network mask or BA44. Comparable results were obtained when computing GLMs across both age groups with children’s syntactic comprehension and production score (see Supplementary Results). In the exploratory analysis reported in the supplementary material, we observed a positive correlation of 4-year-old but not 3-year-old children’s syntactic production scores with their cortical thickness within the mask of BA45 (see Supplementary Figure 3 and Supplementary Table 2). Further, in an additional analysis, we were interested if children’s syntactic comprehension and production abilities show overlapping effects in cortical brain regions within the groups of 3- and 4-year-olds. Within the language-network mask, we found no significant overlap of children’s syntactic comprehension and production scores in neither age group. When performing small-volume correction in BA44, we observed a significant overlap of 4-year-old children’s syntactic comprehension and production scores in this ROI (see Table 1), and no effect for the 3-year-olds.

#### Morpho-syntactic production score

We then tested for the relation of 3- and 4-year-old children’s morpho-syntactic production ability with their cortical brain structure. Analyses within the language network mask yielded a positive correlation of 3-year-olds’ morpho- syntactic production scores with their cortical thickness in the insula (see Table 1), and no effect for the 4-year-olds. Further analysis corroborated that this relation was only present in the 3-year-olds (3-year-olds: *ρ* = 0.78, P < 0.001; 4-year-olds: *ρ* = 0.12, P = 0.63) and that the correlation coefficients differed significantly between age groups (z = 3.94, P < 0.001). The cluster remained significant when controlling for sex, non-verbal IQ and eTIV, but regressed after adding handedness to the model (see Supplementary Material). When testing within BA44, we found a positive correlation between children’s morpho-syntactic production scores and their cortical thickness in this ROI in the 4-year-olds, but no effect for the 3-year-olds (see Table 1). This effect remained significant when controlling for sex, non-verbal IQ, handedness and eTIV. Further analysis corroborated that the effect was only significant in the 4-year-old children (3-year-olds: *ρ* = −0.22, P = 0.40; 4-year-olds: *ρ* = 0.61, P = 0.004) and that the correlations differed significantly between age groups (z = 3.90, P < 0.001). Conjunction analyses yielded no significant overlap between 3- and 4-year-old children’s morpho-syntactic production scores within the language network mask or BA44.

### Discussion

Between 3 and 4 years of age, important behavioral milestones in the acquisition of complex syntax, the foundation for human language, are achieved (Fox and Grodzinsky 1998; Akhtar 1999; Tomasello and Brooks 1999; Guasti 2002; Kauschke 2012). So far, however, little was known about the maturation of the language-related brain network underlying this important achievement in children’s cognitive development. To address this, in the present study, we investigated the association of syntactic abilities with cortical brain structure in 3- and 4-year-old children. Our results suggest that structural maturation of left BA44, a core region for syntax in adults, supports the emergence of complex syntactic structures in 4-year-old children, whereas syntactic abilities in 3-year-olds still rely on the maturation of a different brain region, namely the most posterior left STS, involved in other language or more general processes in adults (Bornkessel et al. 2005; Binder et al. 2009; Friederici et al. 2009; Goucha and Friederici 2015; Bzdok et al. 2016). These effects were not found for children’s global language abilities. These findings suggest a qualitative shift in cortical structures relevant for syntax between 3 and 4 years which may underlie children’s behavioral milestones in this linguistic domain.

The early preschool years are a period of significant, potentially also qualitative, change in syntactic proficiency: By the age of 3, children can reliably process and produce the canonical word order of their language (Akhtar 1999; Dittmar et al. 2008; Schipke et al. 2012), but do not master more complex syntactic structures, such as passive constructions and subordinate clauses, until 4 years of age (Fox and Grodzinsky 1998; Tomasello and Brooks 1999; Guasti 2002; Kauschke 2012). Conversely, a shift from item-based to abstract representation of syntactic constructions between 3 and 4 years has been proposed (Pinker et al. 1987; Akhtar 1999; Bidgood et al. 2020). In the current study, we show that this qualitative change in syntactic development may be reflected in distinct relationships between children’s syntactic abilities and the cortical maturation of different brain regions in 3- compared to 4-year-olds. Particularly, we found that complex syntax mastery in 4-year-olds is supported by the structural maturation of the posterior part of Broca’s area in the left hemisphere, namely BA44. This finding was consistent across all our measures of children’s syntactic abilities, that is, in sentence comprehension and production as well as morpho-syntactic production. In the matured brain, left BA44 is known to support syntactic processes, such as hierarchical structure binding, and shows sensitivity to syntactic complexity and function words conveying structural cues (Friederici et al. 2000, 2006; Friederici 2002, 2011; Makuuchi et al. 2009; Newman et al. 2010; Goucha and Friederici 2015; Zaccarella et al. 2015). Between 4 and 9 years of age, functional activation of BA44 when processing sentences increases, but the neural activity is not yet modulated by syntactic complexity (Skeide et al. 2014, 2016). Our findings are in line with the suggestion that the functional prerequisites for the successful integration of syntax distinct from semantics are already present at that age, but not yet sufficient to master these demanding processing tasks separately (Skeide and Friederici 2016).

In contrast, syntactic proficiency in 3-year-old children was associated with structural maturation in a different region, namely the most posterior left STS. In the matured brain, this brain area has traditionally been associated with semantic processing, as the integration of semantic information and lexical retrieval (Bornkessel et al. 2005; Binder et al. 2009; Friederici et al. 2009; Goucha and Friederici 2015; Bzdok et al. 2016). In the present study, maturation of the most posterior STS was related to the longest syntactically correct fragment that 3-year-old children produced. From behavioral and neuroimaging studies it is known that young children strongly rely on lexical meaning to successfully process complex sentence structures (Friederici 1983; Skeide et al. 2014; Strotseva-Feinschmidt et al. 2019), which is reflected in an immature activation pattern with overlapping neural activity in the posterior part of the superior temporal lobe for syntactic and semantic effects in 4-year-olds (Skeide et al. 2014). The observed association of the most posterior left STS with children’s syntactic abilities may reflect the role of semantic knowledge on the production of longer and thus syntactically more complex utterances in 3-year-olds, as for example they may use a specific sentence construction based on the meaning of a known verb (Pinker et al. 1987; Akhtar 1999; Bidgood et al. 2020). Hence, structural maturation of the most posterior STS may contribute to young children’s ability to construct longer sentences but may not be sufficient for the mastery of complex syntactic structures. The present findings thus fit well with processing and production difficulties of 3-year-olds for more complex sentence structures, such as passive constructions and subordinate clauses, which children typically start mastering from 4 years of age (Fox and Grodzinsky 1998; Akhtar 1999; Tomasello and Brooks 1999; Guasti 2002; Kauschke 2012). While previous studies reported a relation between brain maturation of the left IFG and children’s syntactic comprehension from 5 years on (Fengler et al. 2015; Qi et al. 2019), we showed that structural maturation of BA44 was associated with children’s syntactic abilities in both sentence comprehension and production already at 4 years. However, we did not yet find such a relation at 3 years of age. The qualitative shift from the relevance of the most posterior STS in 3-year-olds to BA44 in 4-year-olds may facilitate the behavioral breakthrough observed in children’s syntax around 4 years.

Despite finding no overlap in the conjunction analysis, we observed a positive correlation of 3- and 4-year-olds’ global language ability and their cortical thickness in the left anterior insula bordering BA44 when aggregating across both age groups (see Supplementary Results). In adults, the left insula is involved in a variety of cognitive processes including working memory demands, language comprehension and production (Flynn 1999; Mutschler et al. 2009; Kurth et al. 2010; Zaccarella and Friederici 2015). The association between structural changes in the insula and children’s global language performance evaluated with a battery of standardized language tests examining children’s linguistic competence across multiple domains fits with findings of its functional involvement in various language tasks in the matured brain. This suggests a gradual development of cortical structure relevant for more general aspects of language ability between 3 and 4 years of age, while the qualitative shift was specific for children’s syntactic abilities.

In addition to our main finding of a dissociated relation between brain regions and syntactic abilities in 3- and 4-year-old children, we also showed that 4-year-old’s syntactic performance in both sentence production and comprehension were associated with maturation of the same region in BA44. Consistent with a recent study in which the authors found broadly overlapping neural activity for sentence production and comprehension in adults (Giglio et al. 2021), our findings indicate similar neural processes involved in the acquisition of both production and comprehension.

In sum, we showed that children’s syntactic abilities were related to maturational changes in different brain regions in 3- compared to 4-year-olds, namely in the most posterior left STS for 3-year-olds, but BA44 in the left IFG for 4-year-olds. This supports a qualitative shift toward more mature brain regions for syntax between 3 and 4 years of age, which may underlie syntactic milestones reached at 4 years. We suggest that the reliance on brain structure of the most posterior STS in 3-year-olds may reflect an influence of semantic knowledge when producing sentences in line with behavioral and functional studies in early childhood (e.g., Friederici 1983; Skeide et al. 2014). To get a better understanding of the role semantics play in sentence production in young preschoolers, future research may investigate children’s processing strategies for syntax before the age of 4 years. The relationship between children’s semantic production and the maturation of brain structures is still poorly understood and may be of interest for further investigation. Moreover, an exciting question for future research is how the left IFG and posterior temporal cortex develops and relates to language abilities and more basic syntactic operations in children younger than 3 years. In behavioral studies, 14- month-old infants for example already show an awareness for syntactic categories suggesting that children’s first knowledge about syntax emerges before the age of 2. Moreover, longitudinal research is required to investigate how brain structure early in life predicts language performance and brain function and structure later on.

## Acknowledgements

We would like to thank Julia Werner, Anne Grigutsch, Lisa Uhlich, and Alina Kowald for their help with correcting segmentation and surface reconstruction in FreeSurfer, and Hung Nguyen Trong and Christiane Attig for their help with data acquisition and organization.

## Author Contributions

**Cheslie C. Klein**: Formal analysis, Writing – Original Draft. **Philipp Berger**: Formal analysis, Writing – Review & Editing. **Tomás Goucha**: Conceptualization. **Angela D. Friederici**: Conceptualization, Writing – Review & Editing. **Charlotte Grosse Wiesmann**: Conceptualization, Formal analysis, Investigation, Writing – Review & Editing.

## Competing Interests

The authors declare no competing interests.

## Notes

### Competing Interest Statement

The authors have declared no competing interest.

