## Supplementary Material for "Children’s syntax is supported by the maturation of BA44 at 4 years, but of the posterior STS at 3 years of age"

Running title: Syntax at 4 but not 3 years relates with BA44 maturation

Cheslie C. Klein[1,2*], Philipp Berger[1,2], Tomás Goucha[1], Angela D. Friederici[1] & Charlotte Grosse Wiesmann[2]

[1]Department of Neuropsychology, Max Planck Institute for Human Cognitive and Brain Sciences, Leipzig, Germany

[2]Research Group Milestones of Early Cognitive Development, Max Planck Institute for Human Cognitive and Brain Sciences, Leipzig, Germany

***Correspondence to:**

Cheslie C. Klein

Max Planck Institute for Human Cognitive and Brain Science

Department of Neuropsychology

Stephanstraße 1a, 04103 Leipzig, Germany

Phone number: +49 341 9940 157

Facsimile number: +49 341 9940 113

### Supplementary Figure 1: Masks for small volume correction

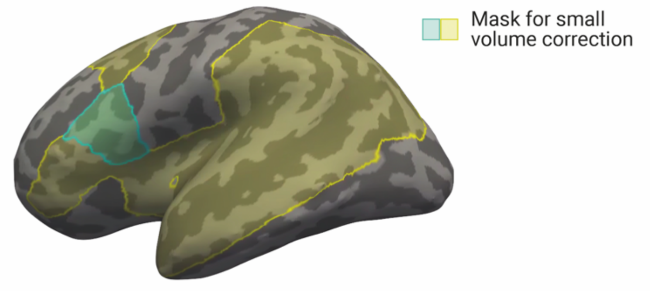


**Figure 1:** Mask of language-related ROIs in the frontotemporal cortex of the left hemisphere using the Desikan-Killiany atlas (light yellow; Desikan et al. 2006) and mask of BA44 for small volume correction (light blue).

### Supplementary Methods: Description of SETK 3-5 tasks

The SETK 3-5 subtest ‘Encoding of semantic relations’ (orig. ‘Enkodierung semantischer Relationen’) is a picture description task in which people and animals perform an action with spatial relation to an object. Therefore, the use of prepositional phrases is elicited in 3-year-olds with varying degrees of difficulty depending on the required preposition (Grimm 1975). In the subtest ‘Sentence memory’ (orig. ‘Satzgedächtnis’, SG), 4-year-olds are asked to reproduce sentences which consist of six to ten words with correct morphosyntactic inflection and either plausible (SG: “Lena lacht, nachdem sie gekitzelt wurde.” [engl. “Lena laughs after being tickled.”]; Grimm 2001) or implausible meaning (SG: “Ein frecher Fußball, der den alten Kasper heiratet, ist müde.” [engl. “A cheeky soccer ball marrying the old Punch is tired.”]; Grimm 2001). The length of the sentences is chosen, such that they cannot be retrieved solely from the working memory of the children but have to be reconstructed using the child’s grammatical and syntactic knowledge (Grimm 2001). This effect is further supported by sentences with implausible meaning, since the children cannot rely on their real-world knowledge (Grimm 2001). The subtest ‘Sentence comprehension’ (orig. ‘Verstehen von Sätzen’, VS) consists of manipulation tasks: The children are instructed, for example, to move objects in a certain order or to touch one object with another. The grammatical complexity of the sentence structure increases from item to item, for instance by presenting the instruction in reverse order than the requested action (VS: "Gib mir die Kiste, nachdem du einen Knopf reingelegt hast." [engl. "Give me the box after you put a button in it."]; Grimm 2001), involving causal relations between two actions and including not only subject- but also object-initial sentences (VS: "Zeig mir: Der gelbe Ball, den der weiße Ball anstößt, fällt vom Tisch." [engl. "Show me: The yellow ball bumped by the white ball falls off the table."]; Grimm 2001). Some items overlap in the tests for 3- and 4-year-old children, but some of the more complex constructions occur only in 4-year-olds as, for example, object-first subordinate clauses are not understood by 3-year-olds (Grimm 2001; Schipke et al. 2012). Instead, the 3-year-olds additionally perform a sentence-picture-matching task in which they are asked to select one of four pictures that matches a sentence. In the subtest 'Morphological rule-formation' (orig. 'Morphologische Regelbildung') children are asked to form the plural of real words. In addition, 4-year-olds are asked to build the plural of pseudowords. The test determines the acquisition level of the morphological rule system for plural formation (Grimm 2001).

**Supplementary Results: Preregistered GLMs with age group as factor**

We had preregistered GLMs including both age groups in one model with age group as factor. Because of task differences between the two age groups based on qualitative changes in syntactic abilities between 3- and 4-year-olds, we decided to run separate analyses in the two age groups instead, and to test for overlapping effects with a conjunction analysis. Following our preregistration, we also ran GLMs across both age groups with age group as factor and tested for an interaction between age group and children's language scores, respectively, within the language-related brain network. In these GLMs, we observed a positive correlation of 3- and 4-year-old children's global language scores with their cortical thickness in a cluster located in the anterior part of the insula extending to BA44 (see Supplementary Figure 2 and

Supplementary Table 1). This cluster remained significant when controlling for age group, sex, non-verbal IQ, handedness and eTIV. Further, we observed a significant interaction between age group and children's global language scores in a cluster located in the posterior part of the left superior temporal gyrus (STG) bordering the Sylvian fissure (see Supplementary Table 1). We then ran separate models within each age group relating children's global language scores with their cortical thickness and surface area. These analyses yielded no significant correlation in the groups of 3- and 4-year-olds. Further, we found no significant cluster and no significant interaction effect for 3- and 4-year-old children's syntactic comprehension scores with a small-volume correction in the language network mask. When testing children's syntactic production scores within the language network mask, we observed no main effect, but a significant interaction with age group in the left posterior STG (see Supplementary Table 1). We then conducted separate analyses of 3- and 4-year-old's syntactic production ability with their cortical brain structure. This revealed a positive correlation of 3-year-old's but not 4-year-olds syntactic production scores and their surface area in the most posterior part of the left STS.

### Supplementary Table 1: Preregistered GLMs with age group as factor

**Table 1:** Mask for small-volume correction (SVC), anatomical region of effect, MNI coordinates, effect size, exact clusterwise P-value and cluster size of significant relation between cortical brain maturation index and language score in the left hemisphere.

| | Mask for SVC | Anatomical region | Peak voxel coordinate in MNI 305 space (X, Y, Z) | Clusterwise P-value | Cluster-size (in $mm^2$) |
|---|---|---|---|---|---|
| **Global language score** | | | | | |
| *Cortical thickness* | | | | | |
| 3y+4y | Language network | Insula, BA44 | -32.4, 4.4, 12.4 | 0.002 | 293.45 |
| *Cortical thickness* | | | | | |
| 3y×4y | Language network | posterior STG | -58.5, -44.2, 26.7 | 0.042 | 187.71 |
| **Syntactic production score** | | | | | |
| *Cortical thickness* | | | | | |
| 3y×4y | Language network | posterior STG | -65.3, -37.6, 12.0 | 0.030 | 200.40 |

Note. Cluster is reported at a cluster-forming threshold of P < 0.01 and clusterwise threshold of P < 0.05.

### Supplementary Figure 2: Global language ability and brain structure across age groups

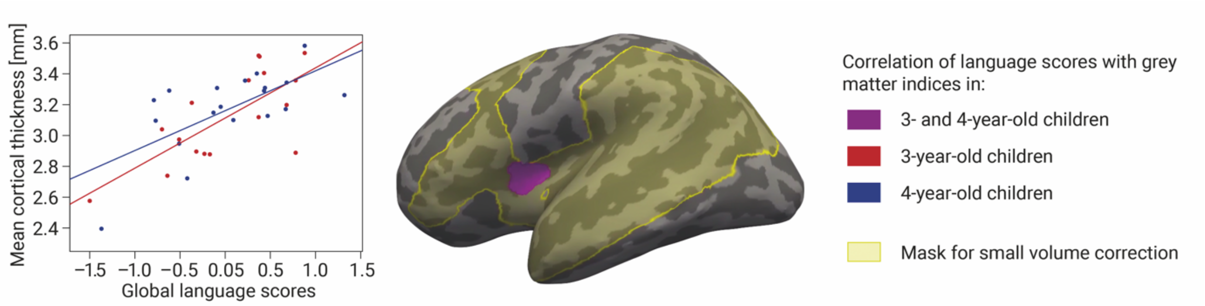


**Figure 2:** Linear correlation of global language abilities of 3- and 4-year-old children (violet) with cortical thickness in the preregistered mask of language-related ROIs (light yellow). The correlation was independent of age group, sex, non-verbal IQ, children's laterality index for handedness and eTIV. The cluster is reported at a cluster-forming threshold of $P < 0.01$ and clusterwise threshold of $P < 0.05$ and displayed on the inflated cortex of the common group template.

### Supplementary Results: Preregistered analysis of the language memory scale in the language-related brain network

We had also preregistered an additional analysis to investigate the relation between children's cortical maturation indices (i.e., cortical thickness and surface area) and their overall performance in the SETK 3-5 subtests assigned to the language memory scale (Grimm 2001). For the 3-year-olds, this scale consists of the subtest 'Phonological working memory for nonwords' (orig. 'Phonologisches Arbeitsgedächtnis für Nichtwörter'), in which children were asked to repeat nonwords of various length. For the 4-year-olds, the subtests 'Memory span for word sequences' (orig. 'Gedächtnisspanne für Wortfolgen'), a repetition task with word lists, and 'Sentence memory' (orig. 'Satzgedächtnis') count in addition to the language memory scale. When performing small-volume correction in the language network mask, no significant effect was found with cortical thickness and surface area in neither age group of 3- or 4-year-olds. Further conjunction analysis to test for overlapping effects in the two age groups revealed no significant relation with cortical thickness or surface area.

**Supplementary Results: Further analyses with right-handed children**

Syntactic production scores of the 4-year-old children were positively correlated with cortical thickness in the left BA44. The effect regressed when controlling for children's laterality index for handedness. To follow up on the effect of handedness, we therefore tested only 4year-olds with a clear indication for right-handedness (n = 14, mean = 71.5, SD = 16.88, range = 42.9 to 100, 7 female). This revealed a significant correlation of children's syntactic production scores with their cortical thickness in the same ROI. The 3-year-old children showed an association between their morpho-syntactic production scores and their cortical thickness in the left insula. The effect regressed when controlling for children's laterality index for handedness. To follow up on the effect of handedness, we therefore tested only 3-year-olds with a clear indication for right-handedness (n = 14, mean = 76.2, SD = 16.01, range = 50 to 100, 9 female). This revealed a significant correlation in the same ROI. The follow-up analyses indicate that these effects were driven by right-handed children.

**Supplementary Results: Exploratory analysis of children's syntactic abilities in BA45**

In an additional analysis, we tested for the relation between children's syntactic abilities and their cortical structures in the left BA45, as this area has been found to be functionally involved in syntactic processing in preschool children and older (Skeide et al. 2014; Wu et al. 2016). No significant correlation of children's syntactic comprehension scores with their cortical thickness or surface area was found in our GLMs in BA45. However, when testing 3- and 4-year-olds' syntactic production scores in this ROI, we found that 4-year-old, but not 3-year-old children showed a positive correlation with their cortical thickness (see Supplementary Figure 3 and Supplementary Table 2). Further analysis corroborated that this relation was only present in the 4-year-olds (3-year-olds: $\rho$ = -0.13, P = 0.62; 4-year-olds: $\rho$ = 0.72, P < 0.001) and that the correlation differed significantly between age groups (z = 4.32, P < 0.001). This effect remained significant when controlling for sex, non-verbal IQ, handedness and eTIV. In the conjunction analyses, we found no significant overlap between 3- and 4-year-old children's syntactic comprehension or production score within BA45.

### Supplementary Table 2: Exploratory analysis of children's syntactic abilities in BA45

**Table 2:** Mask for small-volume correction (SVC), anatomical region of effect, MNI coordinates, effect size, exact clusterwise P-value and cluster size of significant relation between cortical brain maturation index and language score in the left hemisphere.

| | Mask for SVC | Anatomical region | Peak voxel coordinate in MNI 305 space (X, Y, Z) | Clusterwise P-value | Cluster-size (in $mm^2$) |
|---|---|---|---|---|---|
| **Syntactic production score** | | | | | |
| *Cortical thickness* | | | | | |
| 4y | BA45 | BA45 | -40.2, 36.9, -3.8 | 0.025 | 67.48 |

Note. Cluster is reported at a cluster-forming threshold of $P < 0.01$ and clusterwise threshold of $P < 0.05$. It was controlled for sex, non-verbal IQ, handedness and eTIV.

**Supplementary Figure 3: Exploratory analysis of children's syntactic abilities in BA45**

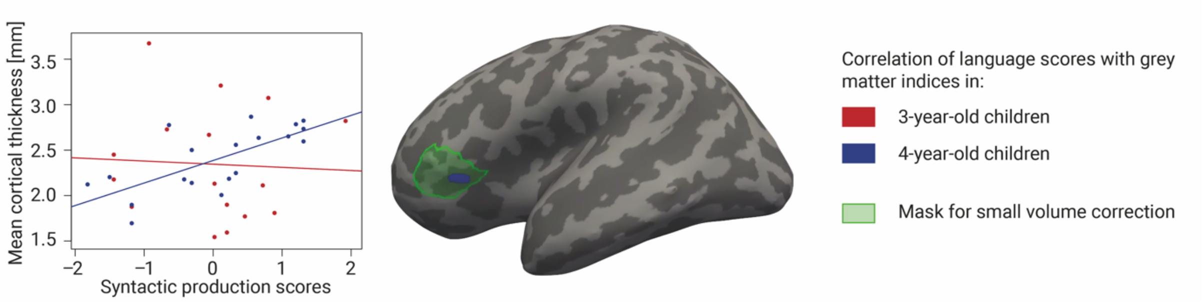


**Figure 3:** Linear correlation of syntactic production abilities of 4-year-old children with cortical thickness in the mask of BA45 (light green). The correlation was independent of sex, non-verbal IQ, children's laterality index for handedness and eTIV. The cluster is reported at a cluster-forming threshold of $P < 0.01$ and clusterwise threshold of $P < 0.05$ and displayed on the inflated cortex of the common group template.
